# Adapting to loss: A normative account of grief

**DOI:** 10.1101/2024.02.06.578702

**Authors:** Zack Dulberg, Rachit Dubey, Jonathan D. Cohen

## Abstract

Grief is a reaction to loss that is observed across human cultures and even in other species. While the particular expressions of grief vary significantly, universal aspects include experiences of emotional pain and frequent remembering of what was lost. Despite its prevalence, and its obvious nature, considering grief from a normative perspective is puzzling: *Why* do we grieve? Why is it *painful*? And why is it sometimes prolonged enough to be clinically impairing? Using the framework of reinforcement learning with memory replay, we offer answers to these questions and suggest, counter-intuitively, that grief may have normative value with respect to reward maximization.

---

Grief is a common reaction to loss, typically including painful recollection of memories and depressed mood (Bonanno and Kaltman, 2001; Weiss, 2008; McCarroll and Yan, 2024). Improving our understanding of grief is important, given not only its prevalence, but also its potential for producing both mental (Nielsen, Christensen, et al., 2020) and physical (Fagundes and Wu, 2020) disability. The study of grief has spanned disciplines and even species (King, 2013). Classical theories of grief include: the “Five Stages" model, describing a sequence of emotional states (Kübler-Ross and Byock, 1969); attachment theory-based descriptions of painful bond breaking (Bowlby, 1979); and the “Dual Process" model, describing oscillation between loss processing and restoration (Schut, 1999). More recently, models informed by neuroscience have suggested that learning (O’connor and Seeley, 2022; Boddez, 2018), representational change (Shear and Shair, 2005), competitive (Békés, Roberts, and Németh, 2023) and cognitive-behavioral (Maccallum and Bryant, 2013) processes play central roles in grief, and may have identifiable neural correlates (Gündel et al., 2003; Seeley et al., 2023).

Despite these efforts, formal computational models that capture the dynamics of grief remain scarce, making it challenging to quantify and predict individual variations in the grieving process. Here, we attempt to address this challenge by leveraging the framework of reinforcement learning (RL) (Sutton and Barto, 2018) to explain the cognitive and emotional shifts occurring during grief. We propose that the psychological pain experienced in grief functions as a form of ‘inverse reward,’ accelerating the update of value representations tied to the object of loss. This mechanism may explain why individuals painfully and repetitively focus on memories of the loss: such memory replay with re-labelled negative rewards serves to expedite the reconfiguration of the value landscape.

To explore this hypothesis, we construct an RL model in which an agent learns about several sources of reward, after which some are removed. We explore how the agent can best respond to this loss. We identify the “grief response" with an operational definition of mood, and vary model parameters to examine how they trade-off with respect to the grief response and subsequent reward. Finally, we compare the behavior of the model to longitudinal data on individual differences in grief reactions, suggesting possible translational applications of our framework to clinical psychology and psychiatry.

## Methods

### Approach

Our hypothesis is that grieving reflects a rational learning process. To formalize this, our work adapts the computational framework of reinforcement learning (RL), and explores the possible learning advantages that grieving provides. RL considers learning in the context of a Markov decision process (MDP), such that at each time-step *t*, an agent perceives the state of the environment *s_t_* and takes an action *a_t_*, causing a transition to state *s_t_*_+1_ and receiving reward *r_t_*. The agent tries to collect as much reward as it can, aiming to maximize the sum of discounted future rewards defined as *G_t_* (Eqn. 1), where γ *∈* [0, 1] is the discount factor, a parameter that scales the present value of future rewards.

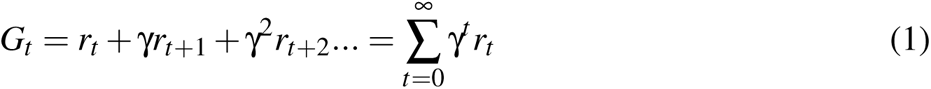

The agent learns to estimate the expected return of action *a* in state *s*, defined as the action value *Q*(*s, a*) (also referred to as the Q-value) as in Eqn. (2).

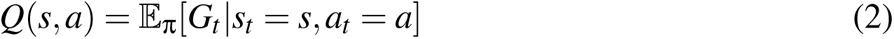

The temporal difference (TD) algorithm updates *Q*-values in state *s* after action *a* transitions to next state *s^′^* (having corresponding action values *Q^′^*), according to learning rate α, as in Eqn. 3 (arguments *s, a* dropped for clarity).

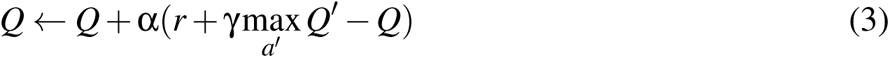

After updating the Q-value of the current state using this algorithm, the agent selects the action with the highest *Q*-value (also known as the greedy action) for the current state; with sufficient learning (i.e., after the agent has accumulated accurate Q-values), this action selection process converges to the optimal behavioral policy (i.e., the one that yields highest cumulative reward). Before this convergence, exploration can assist in updating inaccurately low Q-values, by taking non-greedy actions that would not otherwise be selected. A simple example of this is ε-greedy exploration, during which a random action is selected with probability ε, and the greedy action, argmax*_a_Q*, is selected otherwise (Sutton and Barto, 2018). Typically, the value of ε starts high, and is gradually decreased (annealed) until the optimal policy is learned. Should the agent require additional exploration (e.g., if a change in the environment alters the optimal policy), ε can be increased and annealed again.

TD-learning is known as a model-free strategy (Sutton and Barto, 2018), because value updates depend on current experiences and previously learned values rather than on a model of the environment used to explicitly calculate future values. The gradual learning of cached values from experience in model-free RL takes time and is therefore inflexible if there is a change (like a loss) in the environment; its benefit is that action selection is very efficient. Model-based RL has the opposite trade-off; it can flexibly adapt to changes by simulating, using a world-model, their impact on future reward, but these simulations can be computationally costly. We suggest that, after a loss, grieving reflects the use of a simple model-based mechanism (replaying re-labelled memories; see below) to help increase the efficiency of re-mapping the agent’s model-free value landscape after a loss.

### Problem Set-up

How might an adaptive agent respond to the loss of something valuable? As a first step toward answering this question, we consider a simple version of the problem, involving a *N*x*N* grid-world in which an agent can move in 4 cardinal directions. We label the grid location (0, 0) as *S_A_*(state A), and the location (*N, N*) as *S_B_*. These provide rewards of *r_S_A__* and *r_S_B__*, such that *r_S_A__* (the one that will be lost) is more valuable than *r_S_B__*, with *r* = 0 in all other states (**Fig. 1a**). We first allow the agent to learn the optimal policy in this environment, by taking random actions until its learned action values (Q-values) are stable. Then, *r_S_A__* is set to 0, signifying the loss event. The agent is subsequently moved to *S_A_*, and given *T* time-steps to maximize its reward. The agent must reequilibrate (in analogy to the grieving process, ‘get back to living’) by seeking to collect reward from *S_B_* after the loss of the opportunity for reward from *S_A_*. In the sections that follow, we report simulations that explore the impact of various parameters on the model’s ability to reequilibrate, and maximize its reward, as a formal model of how different strategies may impact the “grieving" process.

**Figure 1.**
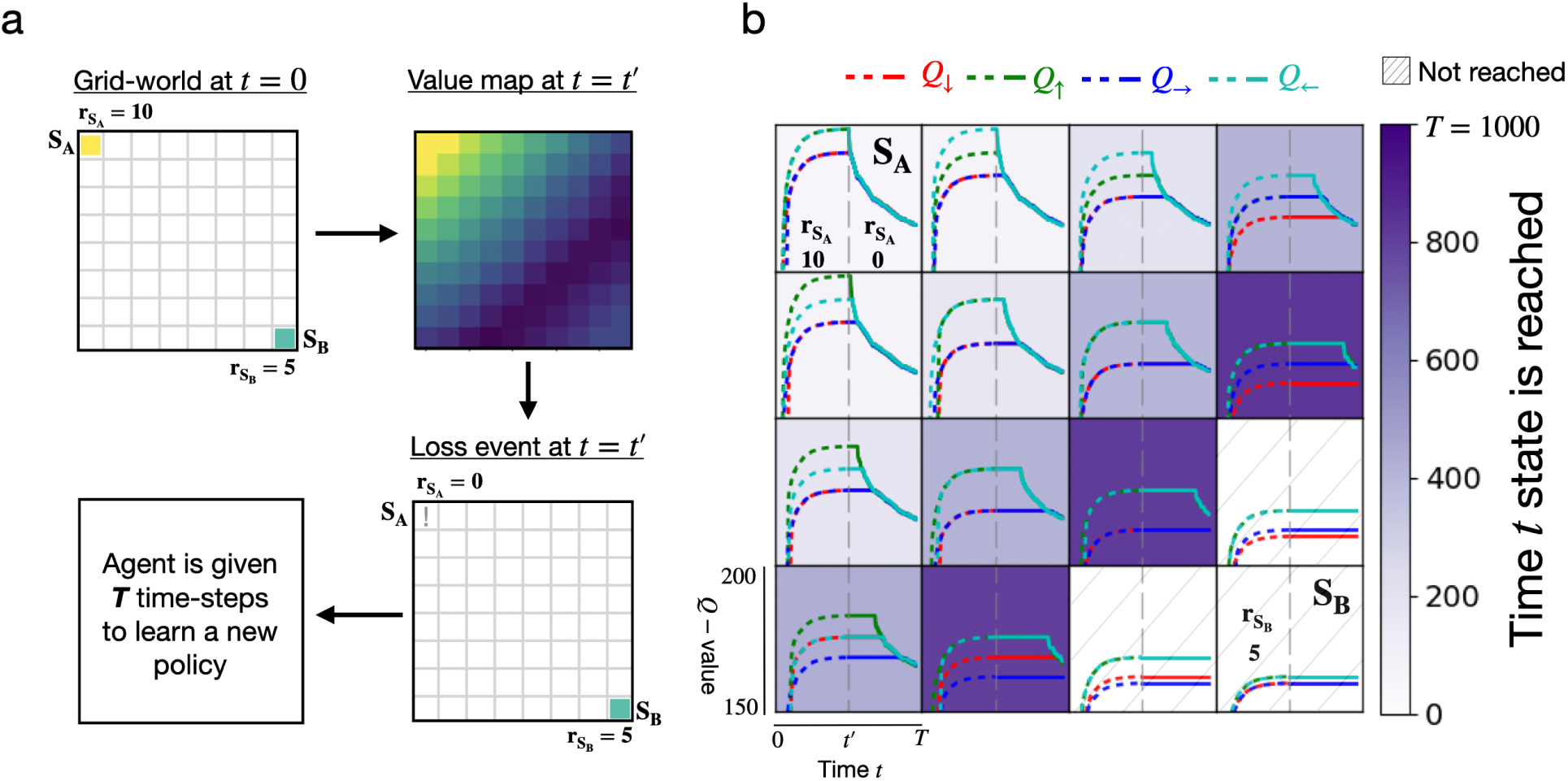
**a: Problem set-up.** Upper left: An 8×8 grid-world in which *S_A_* provides a reward *r_S_A__* of 10 and *S_B_* provides a reward of 5. Upper right: The learned optimal state-value function (i.e. the maximum Q-value in each state after Q-values converge; high value states are brighter.). Lower right: Loss event when the reward provided by *S_A_* is set to 0. Lower left: After the loss, the agent has *T* time-steps to learn a new policy and collect reward from *S_B_*. **b: TD-agent in a simplified** 4**x**4 **grid.** Q-value evolution is displayed over time in each of 16 grid locations of the 4×4 grid (four coloured lines plotted within each grid state, e.g. Q-value of the ‘up’ action *Q_↑_* is in green) before (dashed) and after (solid) the loss event (vertical grey dashed line at *t* = *t^′^*). Additionally, the time at which grid states themselves are first reached after the loss event is indicated in purple shading (darker means later), while hatched states are not reached within an allotted *T* = 1000 steps. The agent starts at *S_A_* after the loss and does not reach the goal state *S_B_*.

### Baseline agent: TD-learning

We begin with a baseline agent using TD-learning (TD-agent), and then explore ways in which its performance can be improved. Our default experiments use an 8×8 grid-world, starting with typical parameter settings: learning rate λ = 0.1, discount factor γ = 0.95, greedy action selection post-loss (i.e. *a* = argmax*_a_Q* with ε = 0), and *T* = 5000 steps to collect reward. One run of this agent is visualized in **Fig. 1b** on a simplified 4×4 grid with *T* = 1000; it never reaches *S_B_*, earning a total reward 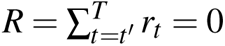 (always calculated post-loss), because model-free TD-learning devalues states slowly as they are experienced, and is unable to adapt in the specified time horizon.

### Grieving agent: Memory replay with re-labelled rewards

How can the process of loss adaptation be accelerated? One obvious way is to increase the learning rate α, and another is to increase exploration (i.e., by increasing initial ε and then annealing it (McClure, Gilzenrat, and J. D. Cohen, 2005; Kaelbling, Littman, and Moore, 1996)). These approaches are reasonably straightforward, but face well known limitations (see Fig. 2a, top panel). Specifically, increasing α is known to cause learning instability and interference (McCloskey and N. J. Cohen, 1989). Similarly, increasing ε can actually worsen performance (since there is no ‘short-cut’ to unlearn the value of *S_A_*, which continues to pull the agent away from *S_B_*, unless ε reaches 0 precisely when the *S_B_* neighborhood is entered).

**Figure 2.**
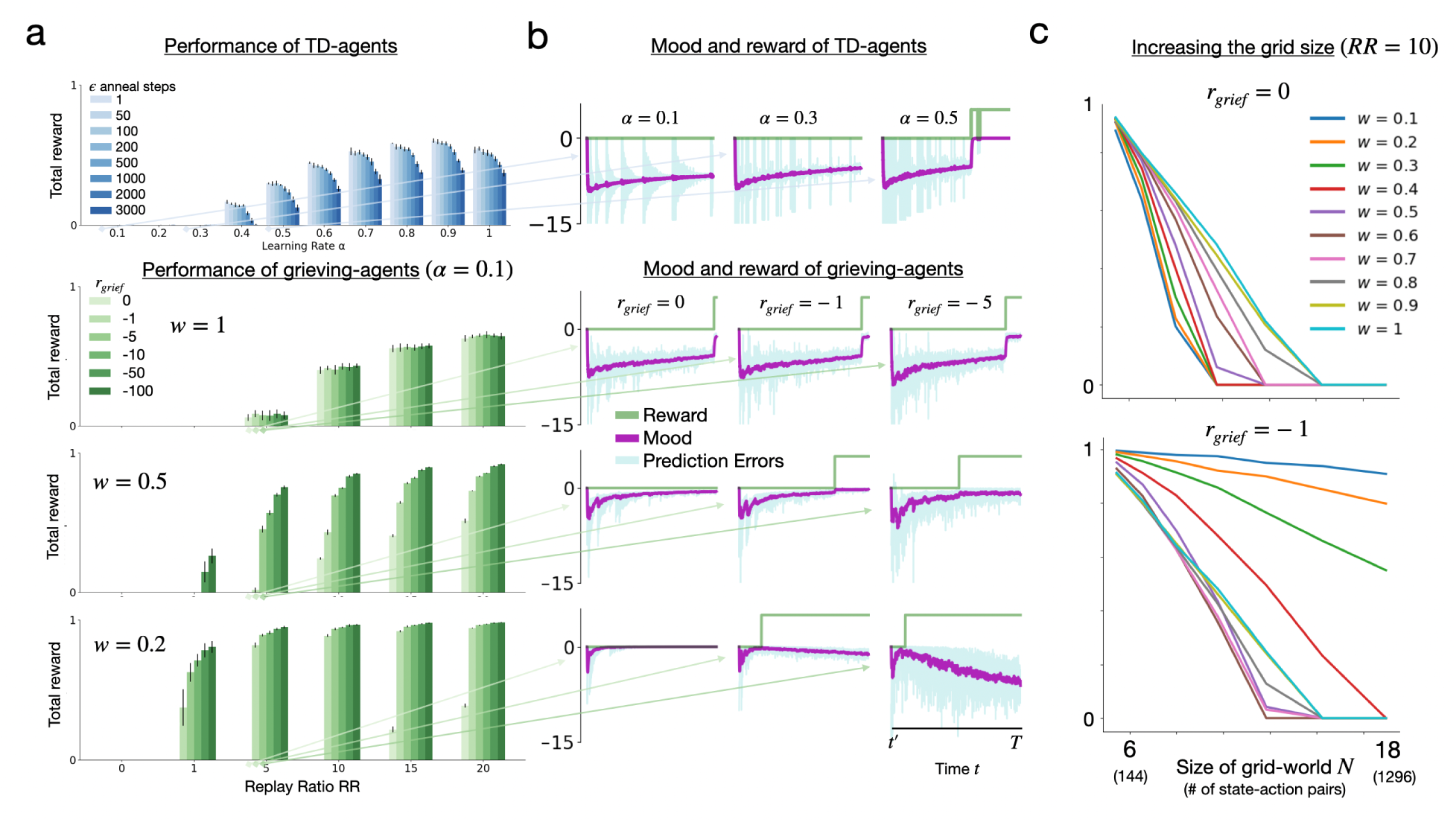
Performance of the TD and grieving agents. **a**: Total reward displayed for the TD-agent over a range of learning rates and annealing steps (blue). For the grieving-agents (green), we set α = 0.1 and plot the total reward for *w ∈* [0.2, 0.5, 1] over a range of replay ratios and *r_grie f_*. Note that the bars show *µ±* σ over 10 runs. **b**: Post-loss trajectory of mood (purple) over *T* = 5000 time-steps (*RR* = 5, η*_replay_* = 0.2, λ = 0.1) and reward (green) with *w* corresponding to adjacent panels in (a). **c**: Total reward of the grieving-agents with increasing grid size *N*, for two *r_grie f_* settings and *RR* = 10. Total reward is normalized so that 1 is the highest achievable score (in both panels a and c).

### Augmenting TD-learning with DYNA, grief, and optimism

A more sophisticated approach is to augment TD-learning with a simple form of model-based learning referred to as DYNA (Sutton and Barto, 2018). DYNA can speed up model-free learning by performing TD-updates using past transitions sampled from a memory buffer (called replay), without having to re-experience them directly. But how should replay proceed after a loss? One way is to use a mechanism, referred to in the machine learning literature as reward relabelling (Andrychowicz et al., 2017; Li, Pinto, and Abbeel, 2020; Eysenbach et al., 2020), in which rewards corresponding to stored state transitions are altered during memory replay. In this case, stored rewards previously experienced in *S_A_*, 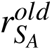, are replayed in the present as 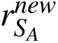 to reflect the loss event.

In the simplest case, old rewards in memory can be substituted with newly experienced ones (i.e. setting 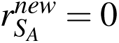 when a reward of 0 is experienced in *S_A_*). Here, we parameterize this choice with a variable called **r_grief_**, such that 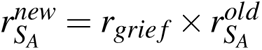. This allows us to accelerate the devaluation of *S_A_* by, for example, setting *r_grie f_ <* 0. By default, we fix *r_grie f_* for the duration of learning after the loss event, reflecting an ongoing process of grief. However, an agent can also stop grieving after some amount of time. To capture this, we also consider a parameter denoted as **t_stop_** (*t_stop_≤ T*), the time point at which *r_grie f_* is re-set to a value *≥* 0.

One problem with reward relabelling in the present setting is that values of *r_grie f_ <* 0 may have little or no effect given the max operator in Eqn. 3, which assumes the best action is always taken, and therefore prevents the value of inferior actions from propagating. However, this is not a realistic assumption; worst-case scenarios matter when actions or state transitions are not deterministic. For instance, even if the edge of a cliff is attractive for the view it provides, its value should be mitigated by the risk of falling off it (e.g., from a gust of wind).

To address this, previous work introduced an *optimism* parameter **w**, which modifies TD-updates (Eqn. 4), maximizing optimism at *w* = 1 (which reduces to Eqn. 3) and maximizing pessimism (i.e., sensitivity to worst-case scenarios) at *w* = 0 (Zorowitz, Momennejad, and Daw, 2020). Critically, the parameter *w* controls the propagation of low values, modulating the impact of decreasing *r_grie f_* in our simulations. We refer to this augmented agent as the **DYNA-***Q_w_***-agent**, that performs updates using Eqn. 4 for both experienced and replayed transitions. In this way, *w* modulates the sensitivity of the agent to worst-case scenarios, because the *minimum* Q-value in a given state increasingly impacts value updates as *w* decreases toward 0.

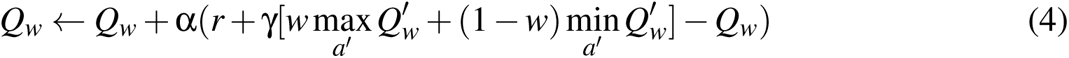

### Deciding which memories to sample

To govern when and how transitions are sampled from memory, we introduce two additional parameters. The first determines the number of transitions replayed per time-step in the environment, commonly called the *replay ratio* and denoted as **RR** (Fedus et al., 2020). Insofar as replay takes time, a high *RR* can be considered a computational cost (Agrawal et al., 2022). The second parameter determines the proportion of memories sampled for replay that pertain to loss-related transitions (i.e., propensity to dwell on grief), that we denote as **p_dwell_**. During memory replay, we sample past transitions in which *S_A_* was the next-state with probability *p_dwell_*, and sample transitions uniformly otherwise.

### Summary

The DYNA-*Q_w_*-agent, that we refer to henceforth as the **grieving-agent**, supplements TD-learning by using Eqn. 4 to update Q-values using ‘painful’ transitions (i.e., re-labelled with *r_grie f_*) sampled from a memory buffer. This is akin to an agent grieving about a previous loss, replaying painful memories that serve to help the agent move on.

### Grief and mood disturbance

Mood disturbance is a major factor in considerations of grief. To explore its relation to grief in our simulations, we follow previous work that has quantified mood by accumulating reward prediction errors (Eldar et al., 2016; Bennett, Davidson, and Niv, 2022); when rewards are consistently lower than expected, the agent is ‘disappointed’ and mood declines, and when they are higher than expected, it is ‘pleasantly surprised’ and mood improves. Eqn. 5 implements this process,

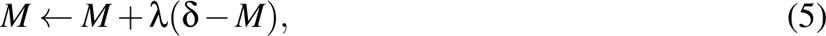

where mood *M* is defined as the running accumulation (at rate λ) of prediction errors δ (Eldar et al., 2016), but with the addition of *w* in the δ term, shown in Eqn. 6 below:

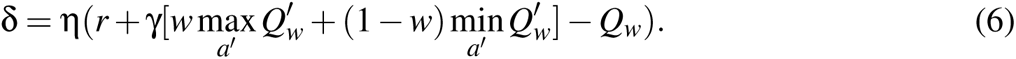

Note that we include the term η, which reflects the differential effect on mood of remembered versus experienced events, by setting it to η*_replay_ ∈* (0, 1) if the transition used to calculate the TD-error was replayed, and (1 *−* η*_replay_*) if the transition was currently experienced.

Finally, we introduce a performance metric *C* (for contentedness) that aims to maximize both total reward *R* and cumulative mood *M* (Eqn. 7), with a relative weighting factor η*_mood_ ∈* (0, 1). Total reward and cumulative mood are normalized to comparable scales by dividing by the maximum achievable reward (see Table S1).

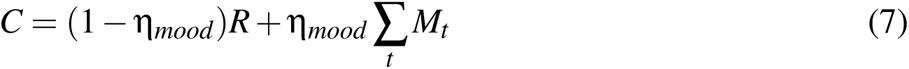

### Overview of Experiments

Using the model described above, we performed a series of simulations to study how grief can improve recovery from loss. To evaluate its performance, we first calculated the total reward for the baseline TD-agent over a range of learning rates α and ε annealing time steps (i.e., the amount of time after the loss event over which ε was gradually decreased from 1 to 0). We then tested the grieving agent over a range of *RR* and *r_grie f_* for different settings of *w*. All agents used ε = 0, either directly after the loss event (grieving-agents) or after annealing was completed (TD-agents). We also calculated the corresponding mood trajectories for a selected set of parameters.

Next, we considered how to rationally optimize the grieving process. For example, how painful should grief be? Given a computational budget (i.e. by fixing the replay ratio), is there a setting of *r_grie f_* that optimizes contentedness, balancing mood and reward? Similarly, how frequently should an agent dwell on their loss? In other words, is there an ideal *p_dwell_* when considering different settings of η*_mood_*? We explored how these and other identified optima (how optimistic should the agent be? How long should grief last?) change as other parameters are varied. In general, all experiments used the default parameter values listed in **Table S1** unless otherwise specified.

Finally, we explored the extent to which variation in *r_grie f_* may help explain clinical syndromes that can sometimes result from grief, such as prolonged grief disorder (PGD) (American Psychiatric Association, 2013).

## Results

### Behavior of learning agents

**Fig. 2a** displays the performance (i.e., the total reward collected after the loss event) of the agents described above. In the top panel, the TD-agent’s performance is shown for different learning rates α (blue plots), and as exploration was increased (shading). We observed that the total reward collected by the agent was 0 until α was increased to at least 0.4, growing gradually until α = 0.9, whereas any amount of time spent exploring hurt performance. Thus, the TD-agent struggled to adapt to the loss.

Typically, RL uses α *≤* 0.1 (and often much lower, depending on how the environment is scaled) to ensure stable learning (Sutton and Barto, 2018). In our simulations, α = 0.1 was also the worst-performing TD-agent tested, and thus provided a useful benchmark against which to compare effects in the grieving agent. Accordingly, the green plots in Fig. 2a show how the grieving-agent enhanced performance with α fixed at 0.1 (and it did so regardless of the specific learning rate). With *w* = 1 (top green plots), increasing *RR* improved total reward, but *r_grie f_*(shading) had no effect. As *w* decreased, lower *r_grie f_* increasingly improved total reward. For example, with *w* = 0.5, *r_grie f_* = *−*10, and *RR* = 5 (middle green plots), total reward was approximately as high as with standard Q-learning with α = 0.9 (Fig. 2a in blue). Further, with more intense and frequent grieving (e.g., *w* = 0.5, *r_grie f_* = *−*100 *RR* = 20), or with lower *w* and only moderate grieving frequency/intensity (e.g., *w* = 0.2, *r_grie f_* = *−*10 *RR* = 10), the total reward was close to its maximum. Thus, the grieving agent was better able to adapt to the loss event, approximating the optimal policy much faster.

Next, we explored the impact of loss on mood. **Fig. 2b** shows the trajectories for mood (purple), raw TD-error (light blue) and reward (green) after the loss event, for a selection of the bars in Fig. 2a (indicated by the faint arrows and labeled above the plots). We found that negative mood appears to oscillate in the grieving agents. For example, this was prominent within the first 1*k* steps of the *w* = 0.5*, r_grie f_* = *−*1 trajectory (third row, middle column). These oscillations emerged as states were successively re-encountered and devalued after loss, and are suggestive of empirically observed ‘waves of grief’ (Zisook and Shear, 2009), even capturing the tendency observed in humans for such waves to decrease in frequency over time (this effect seen most prominently in the light-blue error traces of the TD-agent). In contrast to the grieving agent, the TD-agent progressed through states much more slowly, causing large waves of negative prediction error that resulted in persistently depressed mood; grieving served to compress these waves in time (i.e., increase their frequency) by helping the agent ‘move on’ faster from each state. For the grieving agent, the overall extent of negative mood also depended on model parameters in a non-monotonic fashion (i.e., both high and low *w* induce lower mood than *w* = 0.5 in the context of very low *r_grie f_*). We explore the impact of *w* and other parameters on mood and reward further in the next section.

Lastly, **Fig. 2c** displays performance of grieving agents when the size of the grid-world, *N*, was increased. Specifically, grid dimensions from 6×6 to 18×18 were tested with *RR* = 10, over a range of *w*, and for two settings of *r_grie f_* (with *S_A_*and *S_B_* always occupying top-left and bottom-right corners as in Fig 1). While any decrease in *w* from 1 hurt performance with *r_grie f_* = 0, the previously described interaction between lower *w* and (only slightly) negative *r_grie f_* = *−*1 strongly enhanced robustness to increasing grid-size. This suggests that the grieving model may also have an advantage as environments are scaled up — an important strength since real-world agents must learn in very high-dimensional state-spaces.

### Summary

Augmenting standard TD-agents with grief substantially improved agent adaptation to loss in a simulated grid-world. Specifically, replaying negatively re-labelled loss-related rewards, in the context of an optimism parameter that functioned to increase sensitivity to negative rewards, increased the total reward collected by the agent after a loss. Grieving helped process the loss faster, (a signature of which was increased frequency of mood oscillations), and additionally, enhanced performance robustness to scale as the grid-world dimensions were increased.

### Exploring individual differences: turning parameter dials

To enhance intuition for how model parameters impacted behavior, we visualize how changing parameters one at a time affected agent performance in **Fig. 3**, following the same visualization format as in **Fig. 1b**, but displaying simulations on the larger 8×8 grid-world. Each parameter change inhibited the agent’s ability to adapt in the face of loss, preventing it from reaching the goal state *S_B_* in the allotted time. We also labelled these changes colloquially with potential corresponding clinical / behavioral observations underneath each grid.

**Figure 3.**
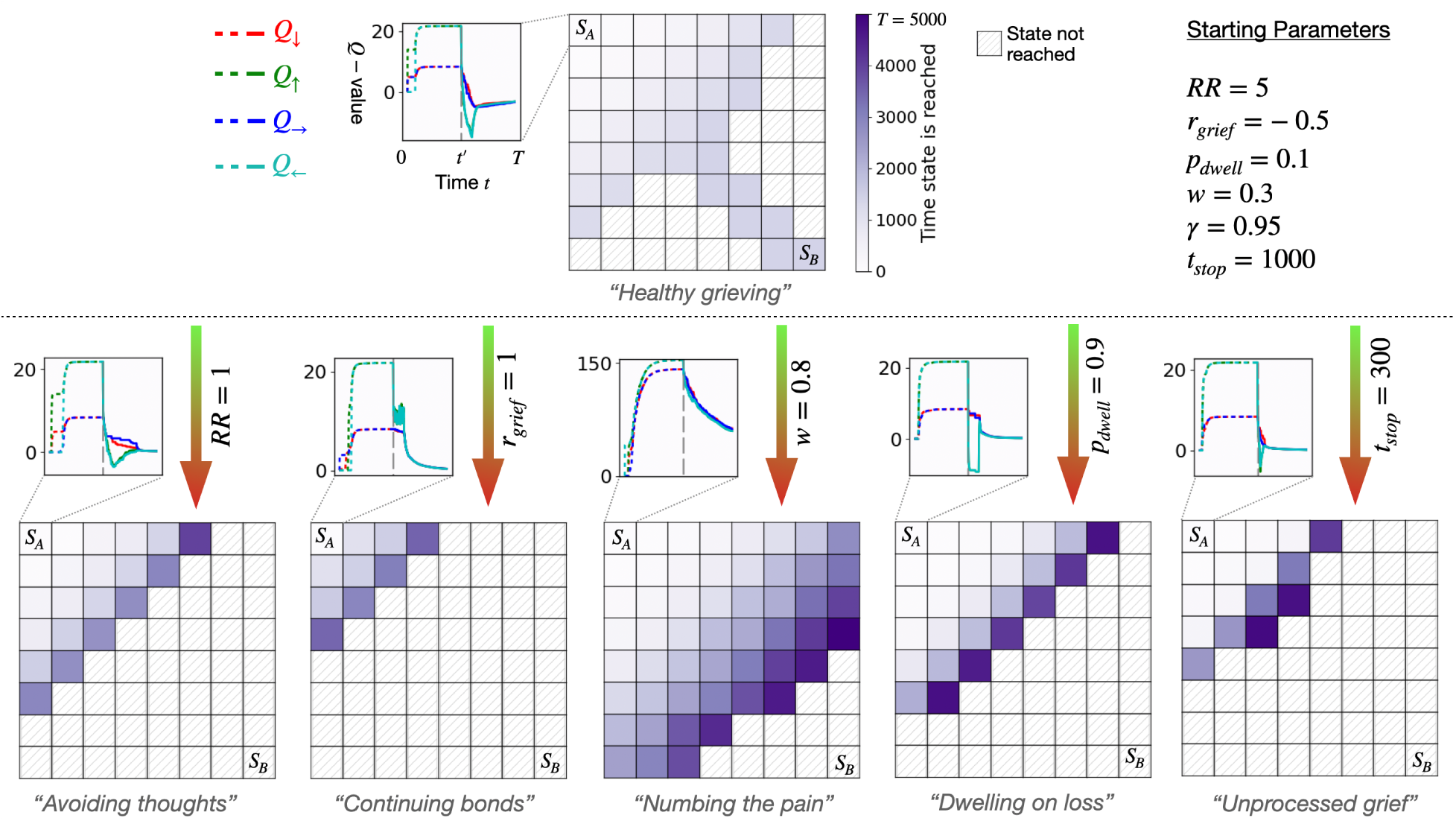
Effect of changing individual parameters on loss adaptation. Adaptation and behavior of agents in 8×8 grid-worlds after a loss is displayed using the same schematic as in Fig. 1b. The agent started in state *S_A_* after the reward in that state was changed to 0 (loss event at *t* = *t^′^*). The agent “recovered" from the loss if it reached goal state *S_B_*. The time at which states were reached after the loss event is indicated in purple (darker means later) and hatched states were not reached within an allotted *T* = 5000 steps. The inset shows the Q-value evolution in state *S_A_*as four coloured traces (e.g., Q-value of the ‘up’ action *Q_↑_* is in green) both before (dashed line) and after (solid line) the loss event (vertical grey dashed line at *t* = *t^′^*). Baseline starting parameters that lead to successful recovery are listed in the top right. Colored arrows indicate a change in one parameter while holding others constant. Each parameter change impaired the recovery of the agent in a different way, displayed as truncated trajectories toward *S_B_*(purple shading) and altered Q-value learning (insets), and also described in colloquial terms underneath each grid.

We began with “healthy grieving", using a set of starting parameters displayed in the upper half of the display. With these parameters, the agent reached *S_B_* within *≈* 1300 steps. The inset displays the learning dynamics of the four Q-values (color coded by action) in state *S_A_*. The loss event occurred at time *t^′^* and is indicated by the vertical dotted grey line. After the loss event, Q-values (solid traces) needed to decrease below their previously learned values (dotted traces) so the agent could move on. We now describe subsequent parameter manipulations in turn.

When the agent performed fewer replays (i.e. setting *RR* = 1 and “avoiding thoughts" about the loss), Q-values decreased less, and the agent did not make it far from *S_A_*. When old memories were replayed in their unaltered form (i.e. setting *R_grie f_* = 1 and “continuing bonds" with the lost object), the agent also stayed close to *S_A_*. In this case, the inset shows oscillation of the maximum Q-value in *S_A_* during grief (light blue; ‘left’ action), since the absence of current reward in experience and the presence of past reward in replayed memories lead to conflicting learning updates. Next, when we decreased agent sensitivity to worst-case values (i.e. setting *w* = 0.8 to “numb the pain" of grief), the agent was again slower to make progress toward *S_B_*. The inset displays how the scale of Q-values were higher and were learned more slowly in this case, given they were driven primarily by maximum values. Continuing on, when the agent predominantly replayed memories involving the lost object (i.e. setting *p_dwell_* = 0.9 and “dwelling in grief"), it again failed to adapt. The inset shows that the value of the lost object decreased fast, however, this did not propagate to other states (not shown), as the agent rarely replayed them, and was therefore unable to widely “integrate" the loss. Finally, when grief was stopped prematurely (i.e. setting *t_stop_* = 300 and leaving behind “unprocessed grief") then learning was not accelerated and the grieving process was not sufficient for the agent to reach *S_B_*.

### Resource rational grieving

While exhaustive exploration of all model parameters is beyond the scope of this work, we examined some informative sets of parameters that revealed optima with potential relevance to psychological phenomena. **Fig. 4** explores points where contentedness *C* — the balance between cumulative reward and mood — was maximized, under the assumption that both reward maximization and mood stability (i.e. avoiding persistent negative mood) were important for the agent (Eldar et al., 2016). In the top row, four parameters were varied (*r_grie f_*, *w*, *p_dwell_*, and *t_stop_*, with other parameters set to default values), and contentedness was plotted for different values of η*_mood_* (green-magenta gradient; η*_mood_* = 0 means only reward matters, η*_mood_* = 1 means only mood matters). The ranges of values over which these parameters were varied were chosen to capture optima that existed in parameter space; one of these optima (i.e., one setting of η*_mood_*) was then selected (light blue arrow), and the parameter value at which the optima occurred, argmax(*C*), was plotted as a function of two additional parameters in the bottom row. The two additional parameters were selected in an exploratory fashion, to check whether optima were robust or varied as other parameters changed. Below, we describe each set of optima in turn, which correspond to separate psychological or behavioral questions.

**Figure 4.**
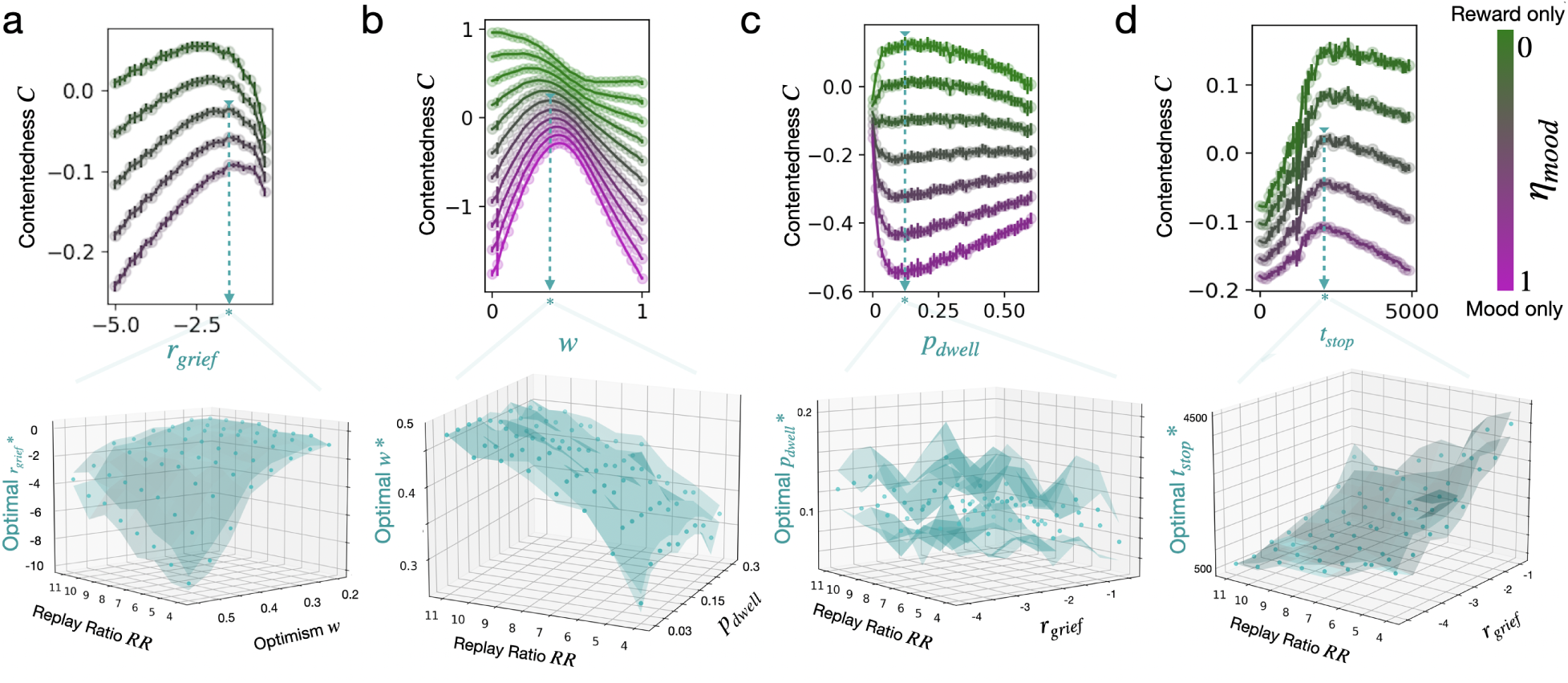
Exploring parameter optima. **Top**: Contentedness *C* as a function of four different grief-related parameters, (**a**) *r_grie f_*, (**b**) *w*, (**c**) *p_dwell_*, and (**d**) *t_stop_*, each for different values of η*_mood_*. Optimal parameter values were computed as the argmax of selected contentedness curves (examples (*) indicated by head of dotted blue arrow). Error bars indicate *µ±* σ over *N* = 10 runs. **Bottom**: Effects of two other parameters on contentedness optima selected in corresponding top panels (indicated by light blue arrow): (**a**) *RR* x *w*, (**b**) *RR* x *p_dwell_*, (**c**) & (**d**) *RR* x *r_grie f_*. Solid points bounded by translucent surfaces indicate *µ±* σ over *N* = 3 runs.

### How painful should grief be?

**Fig. 4a** addresses this question by identifying the optimal value of *r_grie f_*. There was an optimal value because as *r_grie f_* became more negative, more reward was collected, but at the cost of negative mood, and thus these were in tension. The blue surface (in the corresponding panel below) shows how the optimal *r_grie f_* changed as replay ratio *RR* and optimism *w* were varied (considering η*_mood_* = 0.5). A lower *r_grie f_* was needed with lower *RR*, and with higher *w*. That is, with a lower replay budget, or lower sensitivity to negative rewards, a more negative *r_grie f_* was optimal. Thus, to revisit the question about how painful grief should be, Fig. 4a suggests that grief should be more intense when there is a more limited cognitive budget (i.e., lower RR) or when policies are more insensitive to low Q-values (i.e., higher *w*; more optimistic).

### How sensitive should the agent be to the pain of grief?

We address this question by investigating the optimism parameter *w*. In **Fig. 4b**, we found that the extent to which an intermediate value of *w* maximized contentedness was driven entirely by mood: That is, while decreasing *w* (increasing pessimism) monotonically increased total reward (green), both very low and very high *w* negatively impacted mood (magenta). Therefore, a contentedness optima existed where optimism and pessimism were balanced, and was biased toward lower *w* as reward was prioritized. The blue surface shows how the optimal *w* changed as *RR* and *p_dwell_* were varied (considering η*_mood_* = 0.5). Broadly, values of *w* between 0.3 and 0.5 were optimal; as *RR* decreased (lower replay budget), so did the optimal *w*. However, the optimal *w* did not seem to depend on *p_dwell_*. In other words, and to answer our initial question, one should be approximately as sensitive to the pain of grief as they are to positive rewards in general (or more precisely, have similar sensitivity to worst-case and best-case scenarios), but should increase this sensitivity to pain the more limited their cognitive budget. This appeared to be true regardless of the fraction of time spent dwelling in grief.

### How often should the agent dwell on its loss?

We next explore the parameter *p_dwell_*, which reflects the fraction of loss-related memories replayed. The top panel in **Fig. 4c** shows that *p_dwell_* had opposing effects on reward and mood. When reward maximization was prioritized, replayed memories should be related to the loss around 10% of the time (shown by the light blue arrow). However, for an agent that prioritized mood, this was the worst value of *p_dwell_*. In other words, dwelling in grief for the amount of time that was most helpful for value reconfiguration also exacted the greatest cost with respect to mood. If there was too little dwelling in grief, too few prediction errors were accumulated to impact learning or mood. If there was too much dwelling, the newly learned value of the loss state was not propagated to other states, slowing progress toward *S_B_* and also preserving mood by limiting negative prediction errors for more distant states. The blue surface shows how the *p_dwell_* that maximized total reward (i.e., considering η*_mood_*= 0.1) was robust, remaining around 0.1 even as *RR* and *r_grie f_* were varied. Therefore, the answer to our initial question is that agents should dedicate approximately 10% of their time spent remembering the past to grief, but only if they can withstand a parallel cost with respect to their mood.

### How long should grief last?

Finally, we explore *t_stop_*, or the time at which *r_grie f_* was reset to 0. **Fig. 4d** shows that, again there was an optimum: Reward improved if grief lasted until around *t* = 2000, but past that, continued grieving worsened mood without benefit. Naively, we might have expected that this did not occur until the agent reached *S_B_*, at which point there would no longer be a benefit of grieving. However, this is not what was observed (a representative example is shown in **Fig. S5**, in which additional grieving did not improve total reward past *t ≈* 2000, whereas *S_B_*was typically not reached until *t ≈* 3250). The blue surface shows that the optimal *t_stop_* depended strongly on *RR* and *r_grie f_* (considering η*_mood_* = 0.5), decreasing with higher *RR* or lower *r_grie f_*. The answer to our question is therefore relatively straightforward; grieving does not need to last as long if the agent has a higher replay budget, or can tolerate more intense grief, as both would speed up the processing of the loss.

### Summary

In this section, we suggested that grieving is a strategy that maximizes reward, but which comes at a cost to mood, enabling us to calculate the optimal intensity, sensitivity, dwell-time and duration of grief. In other words, we rationally identified the parameters that best supported loss adaptation, given the constraints placed on an agent by resource limitations and by other parameters (Lieder and Griffiths, 2020).

### Accounting for prolonged grief

Prolonged grief disorder (PGD) is defined as a condition in which recovery from grief is impaired and, not surprisingly, is typically associated with unabated emotional distress (Jordan and Litz, 2014). Here, we consider the extent to which the model introduced above may provide insight into how individual differences contribute to the development of PGD. Specifically, we consider how parameters of the model can be used to explore three dimensions of relational attachment style — ‘compulsive care-seeking’ (CCS), ‘compulsive self-reliance’ (CSR), and ‘angry withdrawal’ (AW) — that have been proposed affect the development of prolonged grief (Field and Sundin, 2001).^1^ This study reported that CCS-score was positively correlated with positive loss-related thoughts (*r* = 0.34*, p <* 0.05), and inversely correlated with negative loss-related thoughts (*r* = *−*0.22, n.s.). The opposite was observed for CSR-score, which was negatively correlated with positive loss-related thoughts (*r* = *−*0.53*, p <* 0.001) and positively correlated with negative loss-related thoughts (*r* = 0.36*, p <* 0.05); and for AW-score, which was less negatively correlated with positive loss-related thoughts (*r* = *−*0.35*, p <* 0.05) but more positively correlated with negative loss-related thoughts (*r* = 0.42*, p <* 0.01).

To model these three profiles, we sampled three populations (*N* = 30 agents each) with different *r_grie f_* parameter distributions corresponding to each profile. These were based on the kinds of loss-related thoughts reported to correlate with each attachment style. ^2^ For example, since CCS-score was correlated with positive loss-related thoughts, we simulated a population of CCS-agents by sampling *r_grie f_ ∈* [1, 2]. Similarly, we simulated CSR-agents by sampling *r_grie f_ ∈* [0.1*, −*0.5], and AW-agents by sampling *r_grie f_ ∈* [0.5*, −*1]. Both CSR and AW populations were predominantly comprised of individuals that replayed negative loss-related thoughts, while reflecting that the AW population should consist of agents with more negative *and* more positive loss-related thoughts than the CSR population. We did this by sampling from a wider range of *r_grie f_* magnitudes for the AW population, and also by slightly increasing the frequency of those thoughts, setting *p_dwell_* = 0.05 for the AW population and *p_dwell_* = 0.02 otherwise. **Table 1** summarizes these data and parameters more compactly.

**Table 1.** Human data and model parameters corresponding to different attachment styles.

| Style | Human data |  | Model parameters |  |
| --- | --- | --- | --- | --- |
| | +LRT cor. | -LRT cor. | $r_{grief}$ | $P_{dwell}$ |
| CCS | 0.34* | -0.22 | $\in [1, 2]$ | 0.02 |
| CSR | -0.35* | 0.36* | $\in [0.1, -0.5]$ | 0.02 |
| AW | -0.53*** | 0.42** | $\in [0.5, -1]$ | 0.05 |

\* $p < 0.05$ ; \*\* $p < 0.01$ ; \*\*\* $p < 0.001$ ; two-tailed. Data from (Field and Sundin, 2001).

After sampling three populations of agents, we then simulated the mood time-course of each agent, and used the *r_grie f_* parameter itself as an approximation of the attachment style score. For example, we assumed an agent with *r_grie f_* = 2 would be more likely to have a high CCS-score, while an agent with *r_grie f_* = *−*1 would be more likely to have a high AW-score. We also approximated the level of grief symptoms at a particular time-point by the agent’s current mood at that point, which included the recent history of prediction errors the agent had experienced. Finally, we approximated the correlation within each population between approximated attachment scores and grief symptoms.

**Fig. 5a** shows the empirical correlations measured between grief symptoms and CCS (blue), CSR (magenta) and AW (green) scores over time (Field and Sundin, 2001). As just described, using negative mood *−M* (eqn. 5) as a surrogate for intensity of grief symptoms, and *r_grie f_* as a surrogate of attachment score, we plotted simulated correlations in **Fig. 5b**. While the magnitudes of simulated correlations were higher (presumably due to fewer sources of variance in our model compared to human data), their relative trajectories were very similar to those in the corresponding human populations.

**Figure 5.**
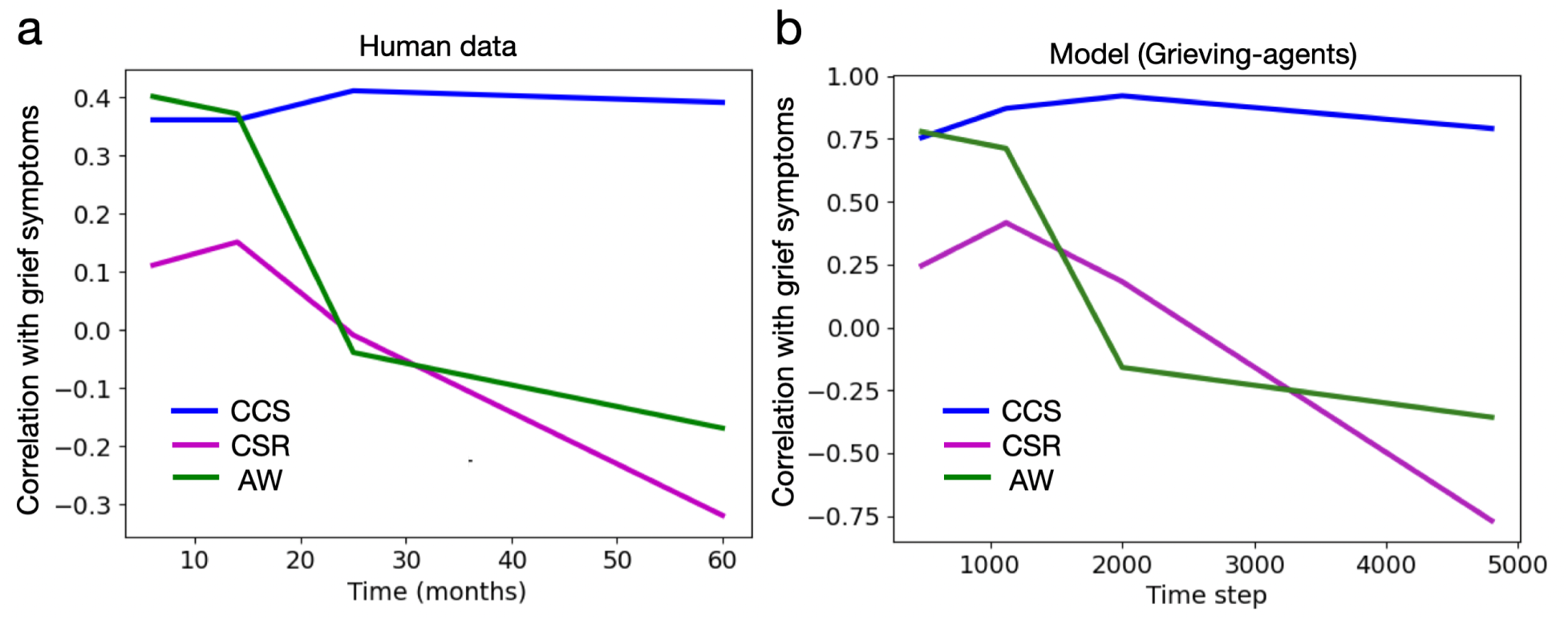
Grief symptom trajectories. **a**: Human data correlating scores on attachment-style dimensions (CCS: compulsive care-giving; CSR: compulsive self-reliance; AW: angry withdrawal) and grief symptom intensity over time (Field and Sundin, 2001). The blue trajectory (CCS) fit the pattern of a prolonged grief reaction, while magenta (CCR) and green (AW) adapted to the loss. **b**: Our model was capable of accounting for the same pattern, defining grief symptoms to be more intense when mood *M* (see eqn. 5) was more negative. The CCS agents replayed positive memories of the loss (*r_grie f_ ∈* [1, 2], *p_dwell_* = 0.02), the CSR agents replayed slightly negative memories (*r_grie f_ ∈* [0.1*, −*0.5], *p_dwell_* = 0.02) while AW agents replayed more negative memories, and to a lesser extent, more positive memories, more frequently (*r_grie f_ ∈* [0.5*, −*1], *p_dwell_* = 0.05). The correlation between *−M* (symptoms) and *r_grie f_* (surrogate for attachment score) was plotted at time steps with the same relative spacing as in **a**. We hypothesized that the valence, intensity and frequency of loss-related memories were the key factors through which attachment style modulates grieving (see main text).

The trajectories in **Fig. 5** along with the data in **Table 1** suggest that replaying positive loss-related memories may prolong grief, while replaying negative ones may shorten it. Our model produced this effect using the following parameters: *RR* = 10, *w* = 0.6, α = 0.1, λ = 0.001 and η*_replay_*= 0. These were close to default parameters used in prior simulations, though it should be noted that it was experienced errors and not replayed errors that produced these particular trajectories (given η*_replay_*= 0; i.e., *r_grie f_* affecting mood indirectly through value propagation rather than directly from painful replays). This is consistent with the possibility that individual differences exist in humans regarding the extent to which memories or experiences drive mood changes in grief.

## Discussion

We showed that augmenting a standard TD-agent with DYNA, *Q_w_*-learning and reward re-labelling improved loss adaptation, and may help explain features of human grieving. Specifically, this work suggests that grief may help to promote the learning of new model-free values after a loss. Grief is painful because, in order to accelerate this learning, the agent must re-experience memories re-labelled with negative rewards. Grief can be prolonged, however, if positive values are maintained for the object of loss (e.g., by the failure to re-label memories). While it has been previously suggested that internally generated negative rewards have computational benefits in accelerating the learning of TD-agents (Dubey, Griffiths, and Dayan, 2022), our work extends this idea to memory replay and reward relabelling, suggesting an additional function of generating negative rewards in the service of loss adaption.

### From model parameters to empirical phenomena: connections and predictions

Computational psychiatry aims to understand how principles underlying mental state dynamics, cast in computationally explicit form, can be used to quantify factors of variation that account for individual differences, and in particular ones that are associated with clinical symptomatology (Huys, Maia, and Frank, 2016). To that end, our results suggest ways in which measurable parameters of learning and replay might be expected to predict grief trajectories.

#### Grief intensity

*r_grie f_*. First, we showed that the lower the time budget for recollection (i.e., low *RR*), and the lower the sensitivity to worst-case Q-values (i.e., high *w*), the more painful grieving should be (i.e., the lower the optimal setting of *r_grie f_*). Existing literature suggests that cognitive fatigue occurs when it is more valuable to replay memories than to continue acting (Agrawal et al., 2022). However, if the cost of replay is high (i.e. because of a limited time budget), an agent would feel fatigued less often. Therefore, our results predict that those who experience less cognitive fatigue may actually experience more intense grief, in order to make the most efficient use of costly memory replay. As well, sensitivity to negative reward has been previously measured in humans (Hundt et al., 2013). Given our results, we would additionally predict that those with a lower sensitivity to negative reward may also experience grief more intensely.

#### Optimism

*w*. Next, with respect to the optimism parameter *w*, we showed that an intermediate-to-low level of optimism was ideal across a range of parameters. Further, a lower value of *w* was optimal as replay budget *RR* decreased, while being relatively robust to changes in the proportion of time dwelling in grief, i.e., *p_dwell_*. This suggests that increasing *sensitivity* to grief is another strategy that could compensate for a limited replay budget (rather than increasing *intensity* of grief as described above). It it unknown which of these parameters would be more controllable, and therefore, unclear which of these strategies humans might prefer.

Additionally, our results suggest that an intermediate-to-low value of *w* would be measured robustly across individuals, which is consistent with the negativity bias (i.e., the tendency to learn from negative more than positive information), that has been observed in humans (Vaish, Grossmann, and Woodward, 2008). It is unclear how *w* might interact with other related constructs that differ between individuals. For example, neuroticism is a personality trait that measures the tendency to experience negative emotion, and those high in this trait are at increased risk of prolonged grief (Boelen et al., 2006). This suggests that the tendency to experience, and sensitivity to, negative emotion may be different constructs, having opposing effects on the efficiency of grieving.

#### Dwell time

*p_dwell_*. We also showed that there is a relatively robust setting of propensity to dwell in grief (i.e., *p_dwell_ ≈* 0.1) that both improves loss adaptation but worsens mood. Therefore, for an agent that prioritizes mood, minimizing grief as well as persistently dwelling in it are both strategies that would avoid negative mood (see Fig. 4c). Consistent with the latter strategy, one study found that the proportion of negative loss-related memories in prolonged grief disorder (PGD) patients was 0.39 *±* 0.27, compared to 0.10 *±* 0.13 in normal grievers (Maccallum and Bryant, 2010). This result in normal grievers is similar to the optimal *p_dwell_*we found in simulation. Corroborating neuroimaging evidence has also shown that PGD patients dwell more in grief-related states of functional connectivity (Seeley et al., 2023). Regardless, either strategy (avoidance or dwelling) would have a negative impact on life restoration, since the *p_dwell_*most effective for value re-mapping would be avoided in both cases by a mood-sensitive agent. Potentially, *p_dwell_* could become a treatment target, predicting that those with a value too low or too high would have a more prolonged course of grief.

#### Grief duration

*t_stop_*. In addition to exploring optimal grieving parameters, we also found that there was an optimal time to stop grieving. This time *t_stop_* likely varies significantly between individuals, given its strong dependence on other parameters such as replay budget and grief intensity in our simulations. Despite this variation, here we briefly explore how *t_stop_* might relate to data on emotional responses during grief trajectories that lead to healthy recovery (regarding when grieving should *start*, see **Fig. S2** for a speculative connection to the phenomena of “anticipatory grief").

Given our hypothesis that grief serves to decrease the value of the object of loss, positive prediction errors might be expected to occur when grieving stops. That is, if *t_stop_* occurs after grieving was sufficient to significantly devalue the object of loss, returning to neutral (or even positive) memories of the loss could induce positive prediction errors (see **Fig. S1** for a simulation of this effect). Related to this, it has been reported that expressions of positive emotion in response to loss-related stimuli six months after a loss contributed to successful recovery from grief (Bonanno and Keltner, 1997). Our findings suggest an interpretation that reverses the causality assumed in this study: positive emotion, rather than promoting recovery from loss itself, is instead a signal that sufficient healthy grieving has already taken place. In other words, positive emotion may be a consequence, not a cause, of successful grieving.

#### Learning strategy

**Model-based vs. model-free.** At the broadest level, our simulations were motivated by the need to update model-free values after a loss. In principle, a fully model-based agent (i.e., one that entirely lacked a model-free system) would not need to grieve lost objects, and could instead simply delete them from its model of the world. Given that individual differences in propensity for model-based vs. model-free decision making can be measured (Daw et al., 2011), and have been shown to correlate with specific psychopathology (Voon et al., 2015), our work also predicts that individuals who rely less on model-free learning would experience less grief.

### From grid to griever: clinical translation in prolonged grief disorder

Lastly, we explored individual differences with respect to clinically-relevant grief symptomatology. We hypothesized that the previously reported modulation of grief trajectories by attachment style (Field and Sundin, 2001) is mediated by differences in the valence, intensity, and frequency of loss-related memory replays. That is, the more positive memories are replayed, the longer the value of the lost object remains high, and the longer grief lasts. Our results comport with somewhat counter-intuitive observations that ‘pleasurable reveries’ can be prominent in PGD (O’Connor and Sussman, 2014), that more calming-feeling memories predict higher grief intensity (Boelen et al., 2006), and that the strongest connection in grief symptom networks is between yearning (i.e., for high-value objects) and emotional pain (Malgaroli, Maccallum, and Bonanno, 2018). They are also consistent with neurobiological evidence that individuals with PGD have higher reward-related neural activity compared to typical grievers (O’Connor, Wellisch, et al., 2008; Bottemanne et al., 2023), as well as higher levels of oxytocin (Bui et al., 2019; Bottemanne et al., 2023), which is known to increase attachment rewards (Scheele et al., 2013).

On this account, replaying positively valenced memories of the lost object maintains and/or increases its value, while the agent consequently experiences negative prediction errors in the post-loss environment - a process that could continue indefinitely. While this likely does not explain all cases of PGD (see Fig. 3 for other hypothesized pathways), it may account for a particular sub-population, for whom novel pharmacological or therapeutic strategies could be suggested. For example, dopamine or opioid receptor blocking agents that target reward circuitry directly could potentially mitigate the preservation of lost object value. In fact, a clinical trial testing Naltrexone (opioid receptor blocker) for the treatment of PGD is already underway; see (Gang et al., 2021). Our model could also inform psychotherapies that precisely target which loss-related memories to recall, and how frequently, in order to optimize value reconfiguration after a loss. The ancient Greek tragedian Aeschylus provided a poetic summary of the present account: *‘There is no pain so great as the memory of joy in present grief’*.

### Future Work

This work can be extended in several directions. First, the parameter *w*, while interpretable, is a heuristic. A more realistic agent would take all Q-values into account, as well as their specific uncertainties, since some actions or transitions may be more deterministic than others. Model-based RL with Bayesian updating or use of successor representations could allow for adaptation to these distributional properties of the environment. Next, memory replay was implemented as a fixed number of random (or loss-biased) samples on each step. It will be important, in future work, to determine whether similar optimal strategies for loss adaptation emerge if the memory replay policy itself is learned (Zha et al., 2019), and if so-called adaptive replay can be extended to accommodate reward relabelling.

Related to adaptive replay, one key component of grief we did not model was avoidance behavior (i.e., avoidance of thinking about painful memories, or avoiding experiences that could trigger them), an additional major driver of PGD (Arizmendi et al., 2023; Eisma and Stroebe, 2021). In our simulations, replayed rewards themselves did not contribute to the agent’s total reward, except as an explanation for why grief is painful. However, interesting avoidance dynamics could emerge if a replay policy had to trade off present painful replays for future reward maximization. In that case, both avoiding painful replays and engaging in pleasurable replays (as previously discussed) could lead to prolonged grief, with their respective contributions potentially accounting for individual differences in PGD.

Another direction would be to incorporate agents with prospective replay (i.e., planning) as well as retrospective replay, since positive counterfactual thoughts about the future is another known contributor to PGD (Golden and Dalgleish, 2012). Future work could also explore when partially retaining the value of lost objects would benefit generalization to related and still-accessible sources of future reward. Achieving a proper balance between retaining and relinquishing so-called ‘continuing-bonds’ is yet another important function of grief (Hewson et al., 2023), and a dimension which would be important to explore in future work.

Lastly, it will be important to extend the results reported here to more ecologically valid environments and reward distributions. First, in complex environments, the impact of a loss may be distributed across many (or infinite) states. This would require multiple different loss-related memories to be first identified, and then replayed to achieve learning that generalizes to future loss-related states. Modular deep Q-learning is a good candidate model for this, since modules maintain different memory buffers corresponding to separate reward components and learn using function approximation (Dulberg et al., 2023). This architecture could therefore naturally accommodate reward re-labelling of replayed memories to generalize learning about the loss of a distinct source of value. Second, given different reward distributions, different emotions would be expected to emerge during loss adaptation. For example, we grieve when we lose valued objects, but are relieved when we lose burdensome ones (see Fig. S3 for a simulation of this effect). Additionally, the same lost object may have contributed a variety of different rewards (e.g., a relationship that had both costs and benefits). Indeed, a mixture of positive and negative emotions are often present in human experiences of loss (Langdon, 2024; Sharbanee and Greenberg, 2023). To capture this complexity, future work could explore loss adaptation in environments with more complex reward landscapes.

### Conclusion

Grief is a fundamental feature of the human condition. In this article, we formalized grief as a solution to the reinforcement learning problem of maximizing reward after experiencing a loss. In constructing a normative basis for grief, we were able to identify optimal grieving parameters, and to reconstruct individual differences in empirically observed human grief trajectories. This model of grief will be a useful foundation for future work in computational psychiatry that aims to further improve our understanding of the complexities of grief, and hopefully, to better predict and treat its manifestations in humans.

## Acknowledgments

Thank you to Isabel Berwian and Simon Segert for helpful discussions. This project / publication was made possible through the support of a grant from the John Templeton Foundation. The opinions expressed in this publication are those of the authors and do not necessarily reflect the views of the John Templeton Foundation. This work was also supported in part by the Office of Naval Research.

## SUPPLEMENTAL INFORMATION

### Capturing additional psychological phenomena

Here, we mention speculative connections that can be made between our model and empirical aspects of a) grief recovery, b) anticipatory grieving and c) relief, and provide simple supporting simulations for each.

### Grief recovery and positive emotion

As mentioned in the main text, expressions of nostalgic-joy 6 months after a loss predicted successful recovery from grief (Bonanno and Keltner, 1997). That is, facial expressions of positive emotion rather than negative emotion when exposed to loss-related stimuli were associated with better subsequent recovery from grief. To explain this phenomena, we first select a set of parameters that should promote optimal grieving based on **Fig. 4** (main text), and observe the resulting mood trajectory. We can see in **Fig. S1**, that when active grieving stops after a sufficient amount of time (i.e. *t_stop_* = 1000, panel **b**), the agent experiences a positive deflection in mood. This larger positive deflection also precedes greater recovery (i.e., greater collection of future reward indicated in green). We hypothesize that the positive emotional expressions observed clinically, which precede and correlate with healthy recovery from grief, reflect this positive mood deflection we observe in our model; a consequence, but not a cause, of healthy grieving.

### Anticipatory grieving

Next, we note that there has been conflicting literature on a phenomena referred to as anticipatory grief (AG), which is when grieving occurs before, or in anticipation of, a loss (Reynolds and Botha, 2006). Classically, AG was reported as positively impacting subsequent bereavement, since some of the work of grief was completed in advance (Lindemann, 1944; Fleming, 1998). There may even be an optimal amount of anticipation, namely, too little (e.g., sudden loss) or too much (e.g., chronic illness) can both lead to more complicated grief reactions (Valdimarsdóttir et al., 2004). In contrast, other studies have reported that AG may increase the risk for complicated/prolonged grief reactions (Nielsen, Neergaard, et al., 2016). Interestingly, pessimism is a mediator of whether anticipatory grief shares features with complicated grief pre-loss (Tomarken et al., 2008). Our model can capture these contradictory findings.

**Figure S1.**
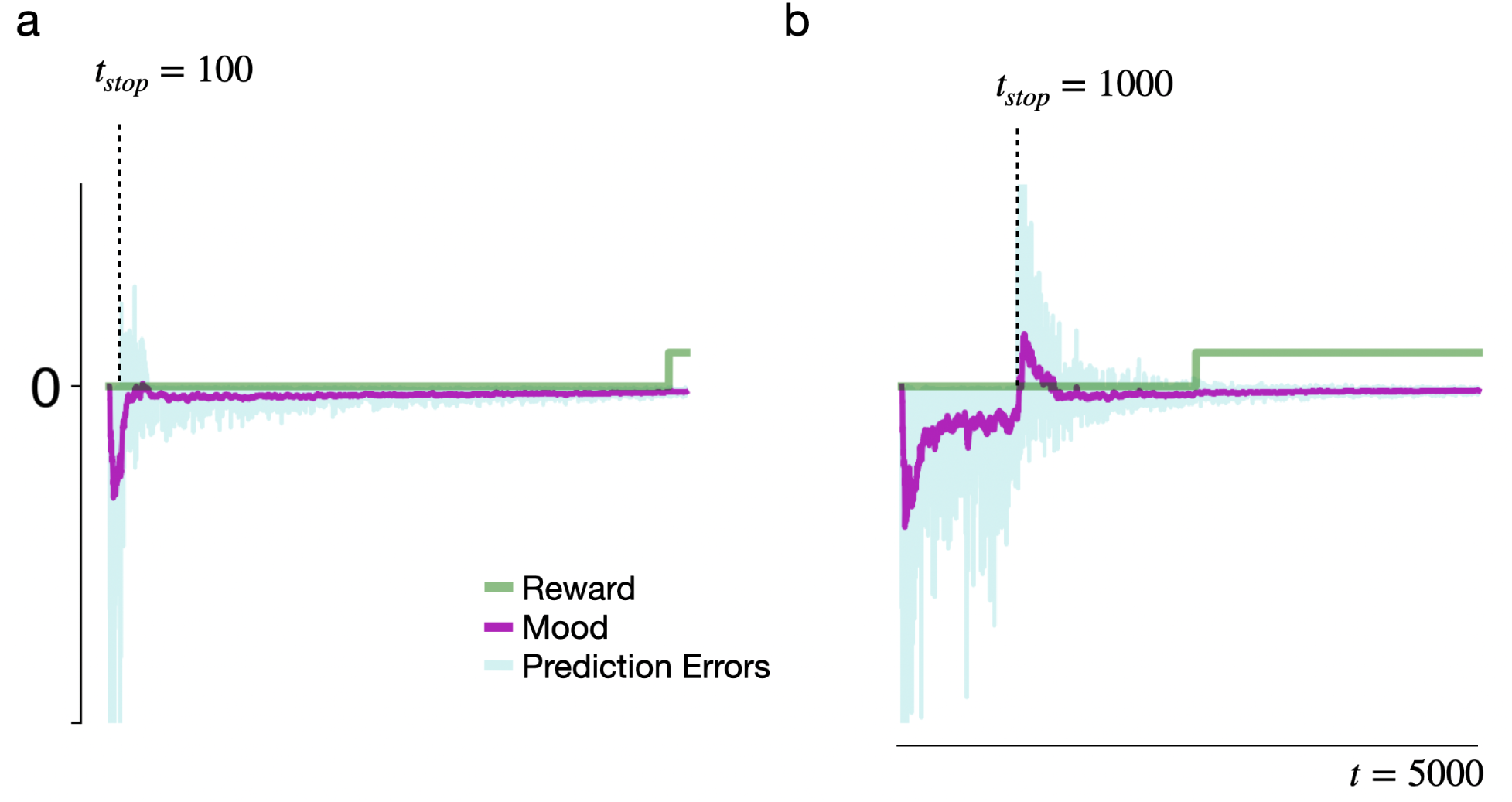
Positive emotion signals healthy recovery. Plots of mood, prediction error, and reward, as in Fig. 2. **a**: Stopping grief early, at *t_stop_* = 100. **b**: Stopping grief later, at *t_stop_* = 1000, closer to the optimal stopping time. When grief stops, the agent experiences a positive deflection in their mood as a consequence of positive prediction errors. The larger deflection in panel **b** also precedes (and therefore would predict) improved recovery from grief (i.e., more reward collected as shown in green) compared with the left panel.

To do so, we introduce another simple parameter called *t_start_*, or the time when grieving starts (when *r_grie f_* is set to a value below 0). Crucially, *t_start_* can be less than *t^′^*, such that grieving starts before the loss. When this happens, the agent experiences conflicting reward signals for the same transition; negative replayed rewards from AG, but positive experienced rewards since the object of value remains present in the environment. As can be seen in **Fig. S2**, this impacts mood in ways that depend on specific model parameters.

The plot shows how cumulative mood and total reward (combined as contentedness *C*) depends on *t_start_*, the time at which grieving begins. As *t_start_* decreases below *t^′^* (occurring from left to right along the x-axis of the graphs), the agent spends more time spent in anticipatory grief before the loss. The top row shows a relatively more optimistic agent (*w* = 0.6) compared with the bottom row (*w* = 0.3). As in the main text, the relative weighting of reward vs. mood is reflected in the green-magenta gradient. On the left, only experienced reward prediction errors contribute to mood, and on the right only replayed ones (indicated by η*_replay_*of 0 and 1, respectively).

**Figure S2.**
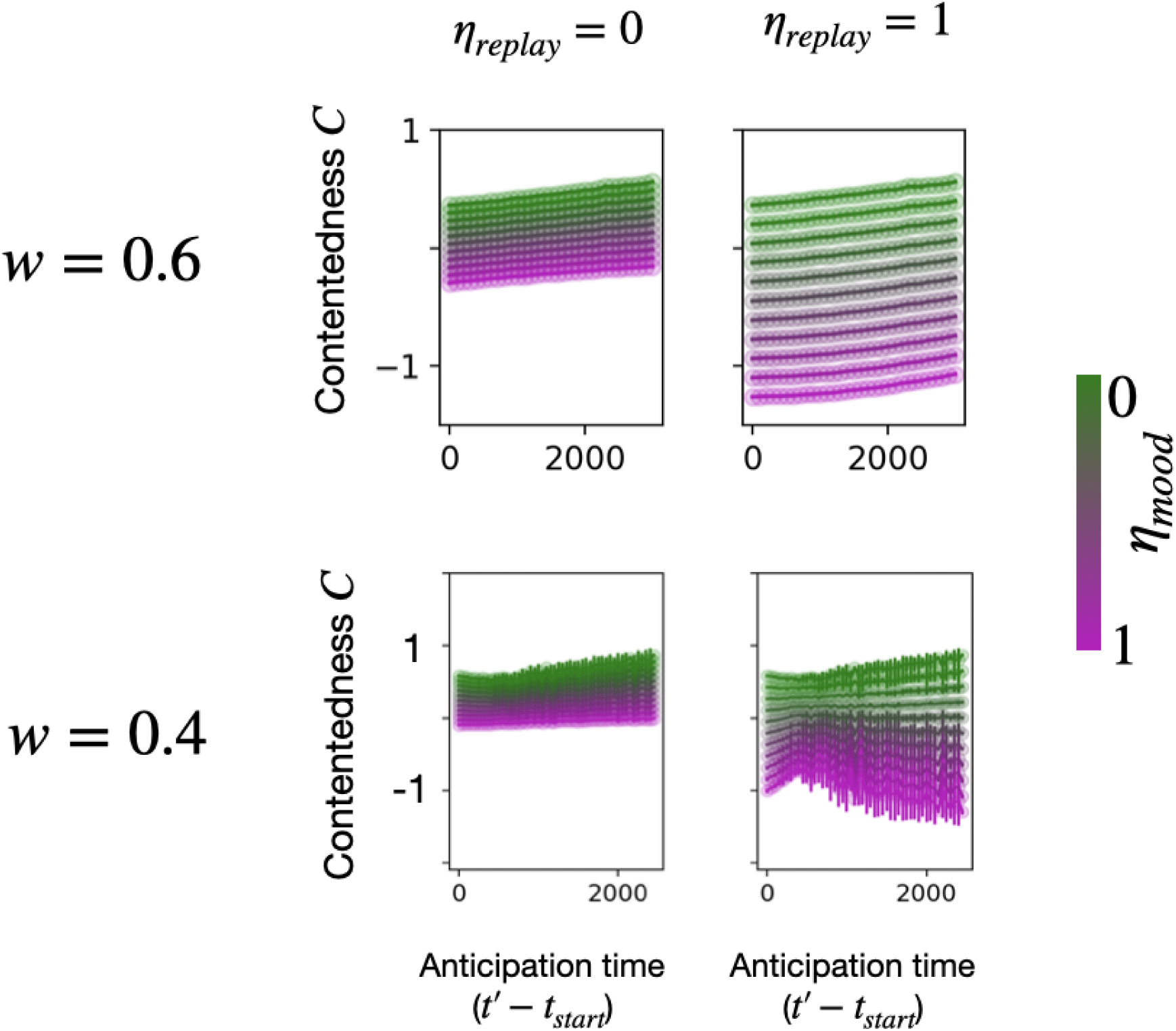
Accounting for anticipatory grief. Contentedness *C* as a function of *t_start_*, the time at which grieving begins. Anticipatory grief occurs when *t_start_ < t^′^* (loss event). The top and bottom rows show relatively more optimistic (*w* = 0.6) and pessimistic (*w* = 0.4) agents, respectively. The relative weighting of reward vs. mood, η*_mood_* is reflected in the green-magenta gradient. The contribution of replayed rather than experienced reward prediction errors on mood, η*_replay_*, is 0 (only experiences) on the left and 1 (only replays) on the right. Error bars show *µ±* σ over 10 runs.

As η*_replay_* increases, a differential effect of *t_start_* on reward and mood develops that depends on *w*. For the higher *w* (optimistic), both reward and mood respond positively to increasing anticipatory grief; as grieving starts earlier (i.e., as *t_start_* decreases), total reward and cumulative mood after loss both increase somewhat, indicating a modest benefit of anticipation. However, for the lower *w* (pessimistic), while anticipation still improves reward, it causes mood to be more negative (i.e. one would expect more grief symptoms). There is even a small optima in mood for the more pessimistic agent around 500 anticipation time steps. This accounts for the contradictory evidence cited above - AG does seem to modestly benefit restoration after loss, could either make mood symptoms better or worse in a way that depends on the pessimism level of the subject, and there does seem to be an optimal anticipation time in the more pessimistic agent.

### Relief: Losing a bad thing

Given the focus of this paper on grief, our modelling assumed the loss of something good; specifically, the state *S_A_* initially provided reward *r_S_A__* = 10, and after the loss provided reward *r_S_A__* = 0. However, it is also possible to lose something bad; life contains many burdens that we may try to avoid. An illustrative example would be if a child experienced bullying, and as a result learned to avoid school. If that bully changed schools, the child may feel relief, and a desire to return to school and capitalize on possible benefits they may have missed out on.

Relief emerges from our model of grief without any substantial changes. Assume that *S_A_* instead started with a reward of *r_S_A__* = *−*5, which changed to *r_S_A__* = 10 after time *t* = *t^′^*. We can again fix a small positive reward in *S_B_* such that *r_S_B__* = 1. In the initial training phase, the agent will learn to avoid *S_A_* in favour of *S_B_*. After *t* = *t^′^*, what should the agent do? A standard TD-agent would remain at *S_B_*, as it would continue to avoid *S_A_* and never learn from new experiences about it. The “grieving" agent, however, would set 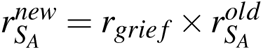 in the exact same way as it did in the main text. Here, again using *r_grie f_ <* 0, but given we also set 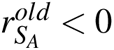, it would replay positively-relabelled loss-related memories. This would help it overcome its avoidance and collect the new source of reward in *S_A_*, again serving to maximize future reward. Both grief and relief are therefore different expressions of the same normative function.

**Fig. S3** shows simulations of total reward and mood in this new environment, just as in main text Fig. 2. As predicted, TD-agents did not perform well (**Fig. S3a**, blue), even as learning rate and exploration was increased. The task was made even easier for the TD-agents, by having them start in *S_A_* after the loss event, whereas grieving agents started in *S_B_*(as far as possible from *S_A_*, the new goal state). Only one TD-agent in 10 runs (α = 0.8, 3000 anneal steps, larger error bar) managed to collect significant reward. For the grieving agents, performance again improved as *RR* increased and *r_grie f_* decreased (**Fig. S3a**, green).

There were two notable differences compared to the main text results. First, high *w*, rather than low *w*, improved performance of the grieving agent. This is because the maximum Q-value was important when the agent had to *increase* the value of the lost state (compared to *decreasing* its value as in grief). Second, as shown in **Fig. S3b**, there is a “wave of relief" that was much larger in magnitude than the waves of grief (and also did not oscillate). This is because of an asymmetry intrinsic to RL: Previously rewarding states are approached, and can be learned about through both experiences and memory replay, while previously punishing states are avoided, and therefore can only be learned about through replay. Therefore, more replay is required to overcome avoidance than to escape attachment.

### Additional baseline model: SARSA

Here, we study an additional baseline agent that uses SARSA learning (which stands for state-action-reward-state-action) rather than Q-learning (Sutton and Barto, 2018). SARSA is an on-policy algorithm, meaning it learns from actually taken actions, rather than best-possible actions, and therefore represents a strategy other than *Q_w_*-learning that also has the potential to propagate low Q-values if the corresponding action is taken. Eqn. 8 shows how SARSA differs from Q-learning (Eqn. 3, main text), where *Q^′^*(*s^′^, a^′^*) is the value of an action actually taken, rather than the action with the maximum Q-value.

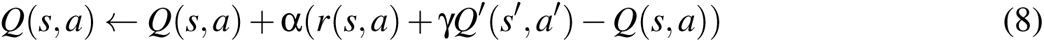

**Figure S3.**
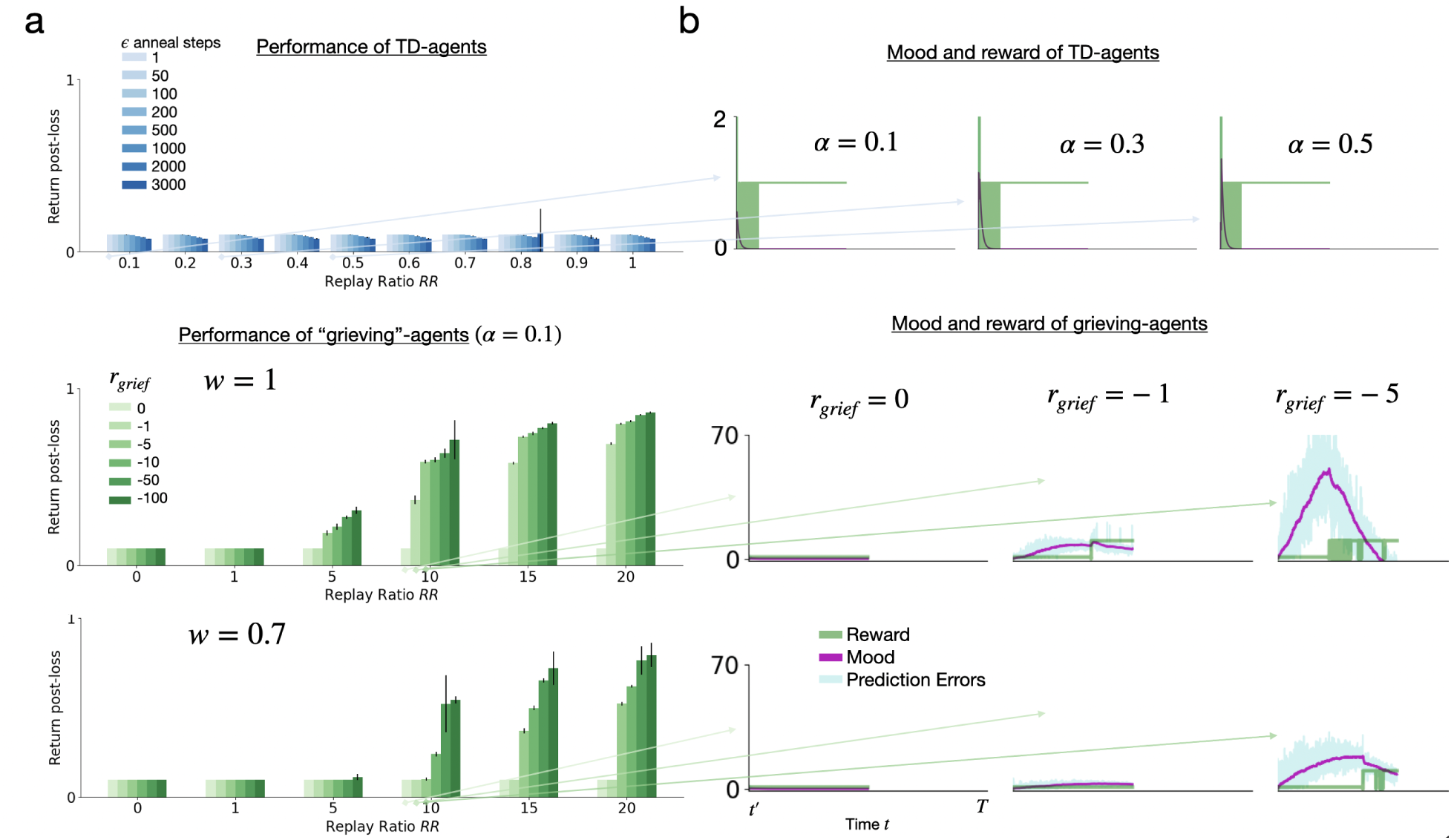
Relief improves adaptation when the object of loss had negative value. Agents were re-analyzed in an environment in which *r_S_A__* changed from *−*5 to 10 at *t* = *t^′^*, and *r_S_B__* = 1 was fixed. **a**: Total reward displayed for the TD-agent over a range of learning rates and annealing steps (blue). For the grieving-agents (green), we set α = 0.1 and plot the total reward for *w ∈* [1, 0.8] over a range of replay ratios and *r_grie f_*. Note that the bars show *µ±* σ over 10 runs. **b**: Post-loss trajectory of mood (purple) over *T* = 5000 time-steps (*RR* = 10, η*_replay_* = 0.2, λ = 0.1) and reward (green) with *w* corresponding to adjacent panels in (a).

Here, we compare SARSA in a setting where memory replay is used, which has previously been referred to as SARSA-DYNA (Al Dabooni and Wunsch, 2016). However, although the task is identical as in the main text (i.e. collect reward after loss with ε = 0), SARSA-DYNA replays memories *as if* ε had been greater than 0 when calculating Q’(s’,a’). The value of this simulated parameter, ε*_SARSA_*, reflects how uncertain the agent is that it can control its actions, i.e., the agent accounts for the fact that if action a was taken from state s into state s’, that a’ could be selected randomly.

**Fig. S4a** displays SARSA-agents compared with a grieving agent (*w* = 0.5) from Fig. 2a (main-text). As predicted, the SARSA-agent is sensitive to *r_grie f_*, improving performance as it becomes more negative. However, even with *e_SARSA_* = 1 (assuming no percieved control over actions), the SARSA-agent still under-performs the grieving agent. This is because its value updates depend on all Q-values, as well as stochasticity in action selection, and not only on the best and worst-case Q-values that the grieving agent uses for updates. This stochasticity is also reflected in noisier mood traces seen in **Fig. S4b**, in which the agent actually experiences a positive mood boost presumably when it reaches a new reward state, and is ‘surprised’ that all of its greedy actions lead to reward, when it had assumed it would lack control over its actions. To what extent humans use Q-learning or SARSA updates is not fully known, and it would be interesting to investigate this in future studies.

**Figure S4.**
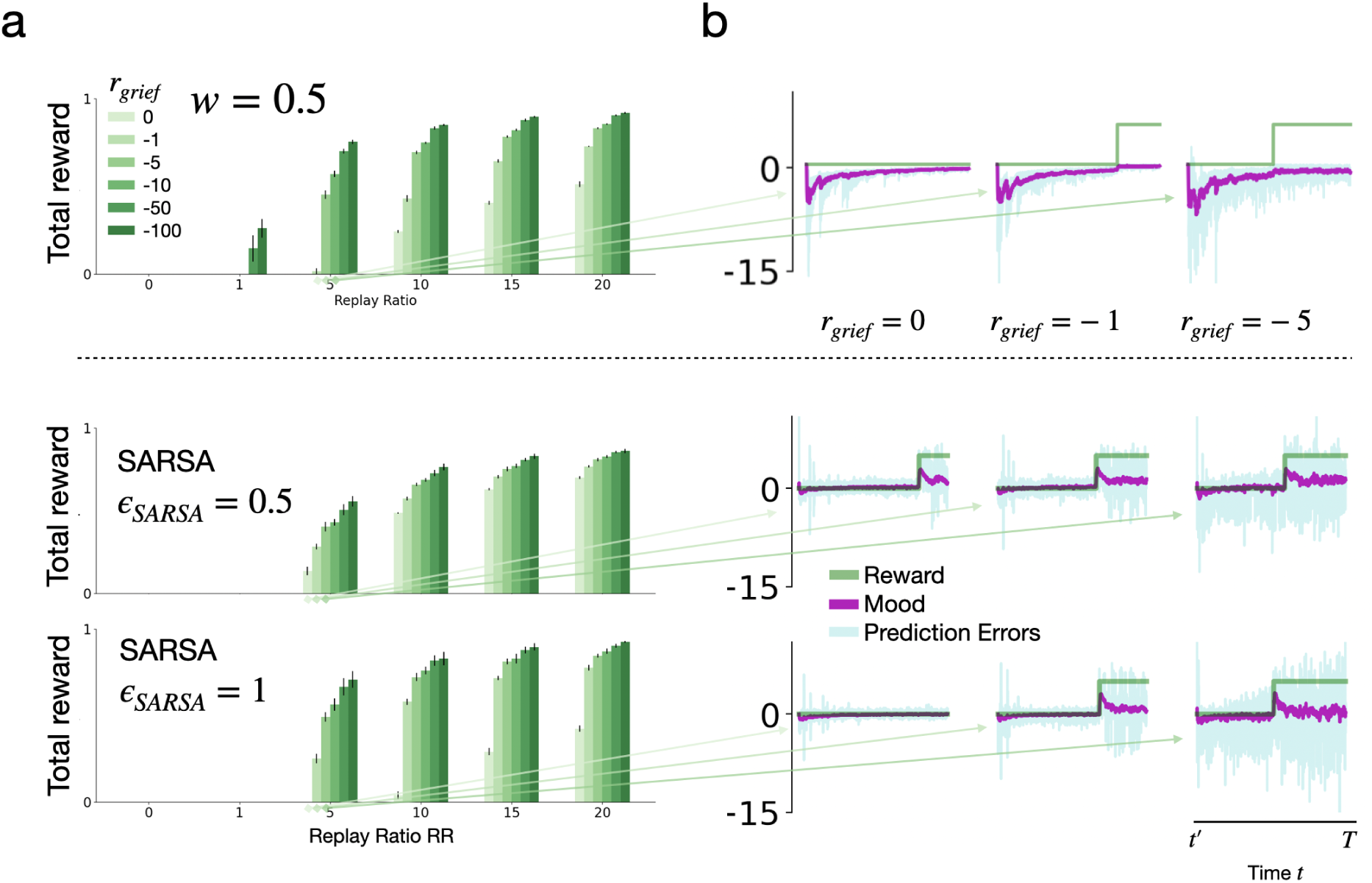
Performance of SARSA agents. **a**: Performance of grieving-agents with *w* = 0.5 (top), compared to SARSA agents (bottom) for two settings of ε*_SARSA_*. We set α = 0.1 and plot the total reward over a range of replay ratios and *r_grie f_*. Bars show *µ±* σ over 10 runs. **b**: Post-loss trajectory of mood (purple) over *T* = 5000 time-steps (*RR* = 5, η*_replay_* = 0.2, λ = 0.1) and reward (green) corresponding to adjacent panels in (a).

### Additional figures

**Figure S5.**
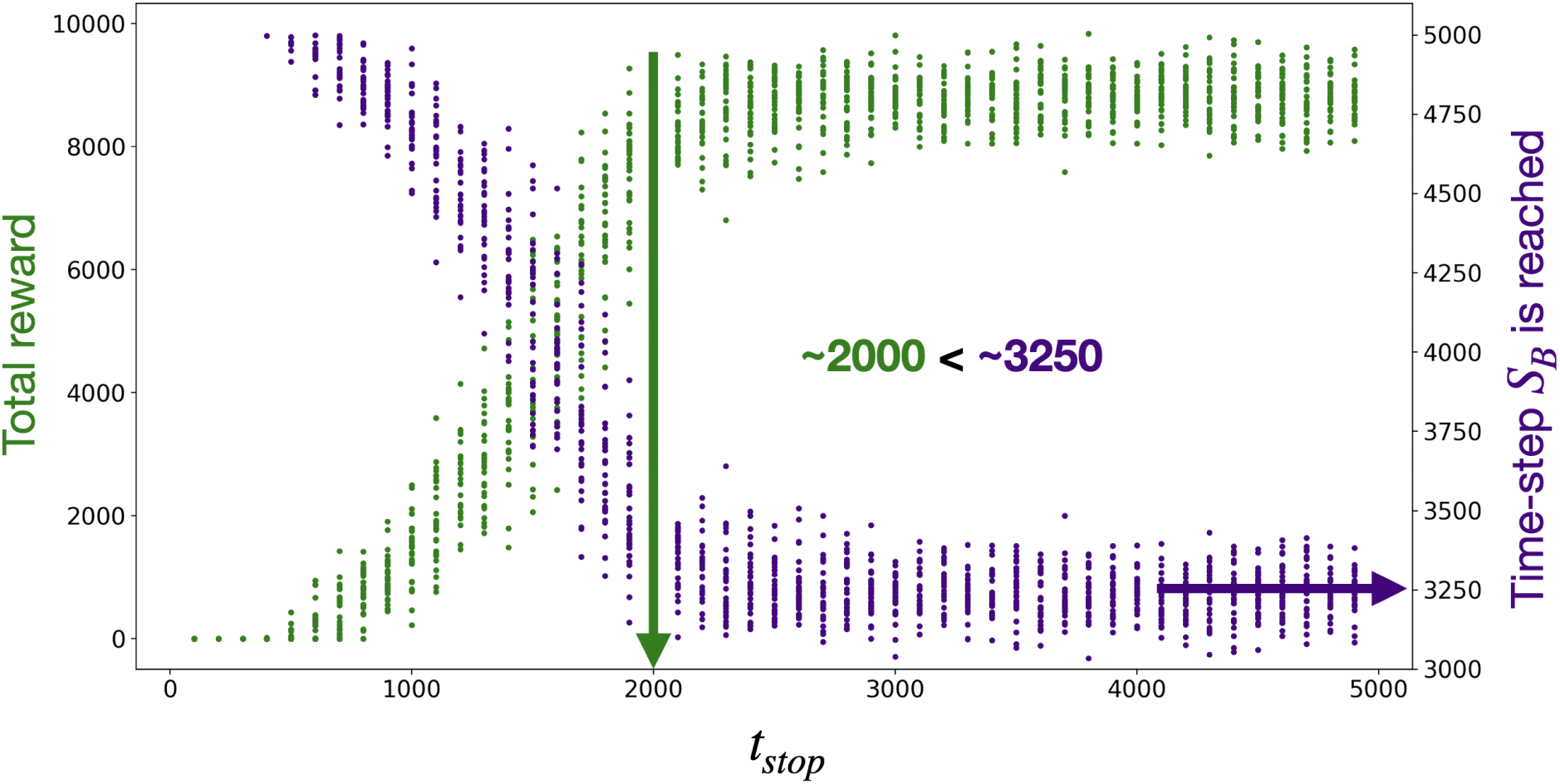
Grief should stop before *S_B_* is reached. In green, we plot the total reward as duration of grief, *t_stop_*, is increased (here, the loss event occurs at *t* = 0). Each point shows a single agent; 20 agents were run for each value of *t_stop_*. With little or no grieving, i.e. *t_stop_ ≈* 0, total reward is 0, gradually going up as *t_stop_* increases, but reaching plateau around *t_stop_ ≈* 2000. In purple, we plot the time step after the loss event when the goal state, *S_B_*, is first reached. One can see that *S_B_* is never reached earlier than *t_stop_≈* 3250. That is, the point past which there is no longer any benefit to further grieving is not the trivial point when *S_B_* is reached; it occurs much earlier in time.

### Default Parameters

**Table S1.**
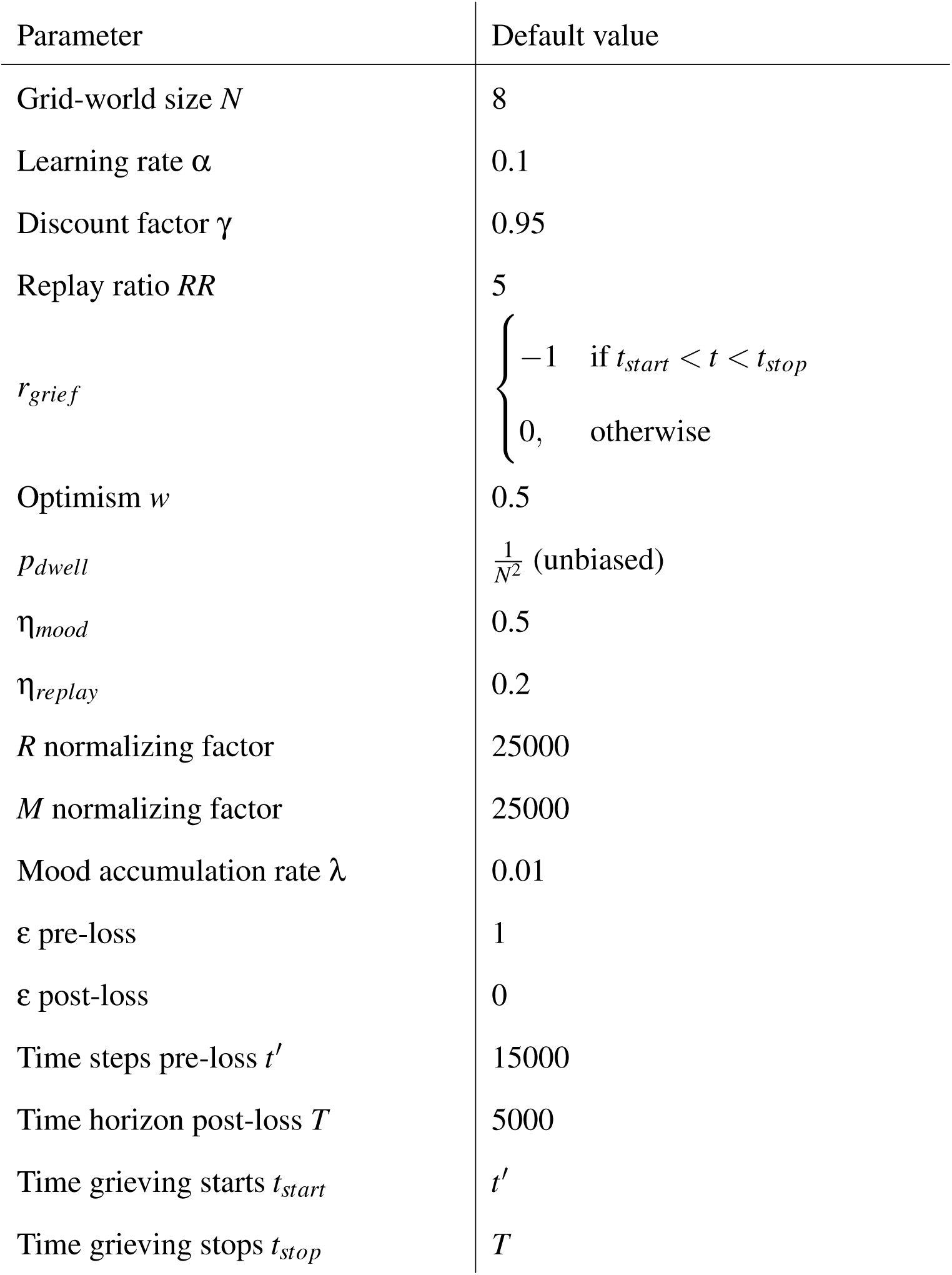
Default parameters used in all simulations if not otherwise specified.

## Footnotes

1 While Field & Sundin reported these results over a decade before Prolonged Grief Disorder (PGD) was introduced in the Diagnostic and Statistical Manual of Mental Disorders, Fifth Edition (DSM-V), we refer to the phenomena of prolonged/complicated grief and the formal disorder PGD interchangeably.

2 Field & Sundin reported that “the extent of positive and negative thoughts and feelings about the deceased" was captured on a numerical scale from 1 (not at all) to 5 (very much) (Field and Sundin, 2001). However, questionnaire items were not reported, so the way in which they captured the magnitude and/or frequency of thoughts had to be assumed in our modeling.

